# Reference-free variant calling with local graph construction with ska lo (SKA)

**DOI:** 10.1101/2024.10.02.616334

**Authors:** Romain Derelle, Kieran Madon, Joel Hellewell, Víctor Rodríguez-Bouza, Nimalan Arinaminpathy, Ajit Lalvani, Nicholas J. Croucher, Simon R. Harris, John A. Lees, Leonid Chindelevitch

## Abstract

The study of genomic variants is increasingly important for public health surveillance of pathogens. Traditional variant calling methods from whole-genome sequencing data rely on reference-based alignment, which can introduce biases and require significant computational resources. Alignment-free and reference-free approaches offer an alternative by leveraging k-mer-based methods, but existing implementations often suffer from sensitivity limitations, particularly in high mutation density genomic regions. Here, we present ska lo, a graph-based algorithm that aims to identify variants between pathogen whole-genome sequencing data by traversing a coloured De Bruijn graph and building variant groups (ie, sets of variant combinations). Through in-silico benchmarking and real-world dataset analyses, we demonstrate that ska lo achieves high sensitivity in SNP calls while also enabling the detection of insertions and deletions, as well as SNP positioning on a reference genome for recombination analyses. These findings highlight ska lo as a simple, fast and effective tool for pathogen genomic epidemiology, extending the range of reference-free variant calling approaches. ska lo is freely available as part of the SKA program (https://github.com/bacpop/ska.rust).

## Introduction

The study of genomic variations plays a central role in public health surveillance, enabling scientists to trace the evolution of pathogens, predict antimicrobial resistance, and monitor the spread of infectious diseases (Simon R. Harris et al. 2010; Quick et al. 2016; Armstrong et al. 2019). These insights are critical for designing effective interventions and controlling outbreaks as demonstrated by the recent COVID-19 pandemic (Cyranoski 2021; Stockdale et al. 2023). Traditionally, the detection of genomic variants from whole-genome sequencing (WGS) data relies on aligning sequencing reads to a reference genome. While effective, its accuracy is highly dependent on the choice of the reference genome (Stephen J. Bush et al. 2020; Valiente-Mullor et al. 2021; Colquhoun et al. 2021), and is controlled by a large set of threshold parameters that require technical expertise to be optimised and, in some cases, species-specific customization (S. J. Bush 2021). Despite the declining costs of WGS, the technical complexity and computational cost of bioinformatic approaches such as read-alignment have limited the expansion of genomic surveillance programs, particularly in resource-limited settings where it is needed most.

Alignment-free and reference-free variant-calling approaches emerged over a decade ago as alternatives to traditional reference-based pipelines, offering simple and fast solutions for analysing WGS data but have so far failed to gain significant traction due to their lower sensitivity (but see Discussion). An alignment-free algorithm typically consists of reducing WGS data to k-mers and storing them in data structures that allow direct variant calling. One of these data structures is the split k-mer, whereby k-mers are subdivided into two smaller k-mers surrounding a middle base (S. R. Harris 2018). Single nucleotide polymorphisms (SNPs) are then called by comparing the middle-bases of surrounding k-mers shared by different samples, allowing ultra-fast SNP identification in reference-free mode. This approach has been implemented in the programs kSNP (Hall and Nisbet 2023; Gardner and Hall 2013), DiscoSNP (Uricaru et al. 2015) and SKA (command ska align) (Derelle et al. 2024; S. R. Harris 2018). However, its sensitivity depends on the conservation of surrounding k-mers between samples, which can be compromised by the presence of multiple SNPs within k nucleotides in a single sample (multiple nucleotide polymorphism, hereafter abbreviated to MNP) or variants occurring within k nucleotides across samples (hereafter referred to as ‘overlapping variants’). For these reasons, split k-mer analyses are typically limited to closely related genomes (S. R. Harris 2018; Moore et al. 2024; Derelle et al. 2024). Alternatively, the sets of split k-mers in each sample can be mapped one by one onto a reference genome to avoid the loss of SNPs associated with overlapping variants (command ska map). The second main data structure is the assembly graph, in which overlapping k-mers form a directed graph that is traversed to identify bubbles. Variants, including SNPs and others, are called by comparing the paths within each bubble, as implemented for example in Cortex (Iqbal, Turner, and McVean 2013), DiscoSNP++ (Peterlongo et al. 2017), Kmer2SNP (Li, Patel, and Lin 2022), MALVA (Denti et al. 2019) and KAGE (Grytten, Dagestad Rand, and Sandve 2022), all primarily designed for analyses of diploid genomes. While this approach enables the detection of MNPs, it theoretically remains susceptible to sensitivity loss due to overlapping variants, as only samples sharing k-mers with the bubble paths are typed for the corresponding SNPs.

Notably, both of these reference-free data structures share a fundamental similarity in the variant calling process: in split k-mer analysis, the two surrounding k-mers serve as local reference points for SNP detection, just as the entry and exit nodes of graph bubbles function as anchors for identifying variants in assembly graphs. Here, we introduce ska lo, a new command of the SKA toolkit (Derelle et al. 2024), that extends the assembly graph a step further by considering all variant paths located between any given set of entry and exit nodes (hereafter referred to as ‘variant groups’). In doing so, ska lo is able to recover nucleotides in samples that are not part of individual SNP bubbles and is therefore less affected by overlapping variants in highly polymorphic regions. SNPs are also extracted from MNPs but with some limitations. Finally, it can position SNPs on a reference genome to conduct recombination analyses. ska lo also identifies insertions and deletions (‘indels’) using graph bubbles as structural markers to detect these variations. (Derelle et al. 2024). In this study we provide a summary of the algorithm, discuss its limitations, and evaluate its performance using in-silico benchmarks and real-world datasets, including outbreak and within-strain sequencing datasets.

## Materials and methods

### Algorithm description

ska lo is a multi-core command of the SKA program (SKA v0.4; https://github.com/bacpop/ska.rust), whose name stands for ‘SKA left over’. As with other SNP-calling SKA commands, it takes as input a multi-sample split k-mer file, which contains all split k-mers of size ‘k’ extracted from assemblies or read data. Its algorithm can be divided into five steps, as summarized in Figure 1:

**Figure 1:**
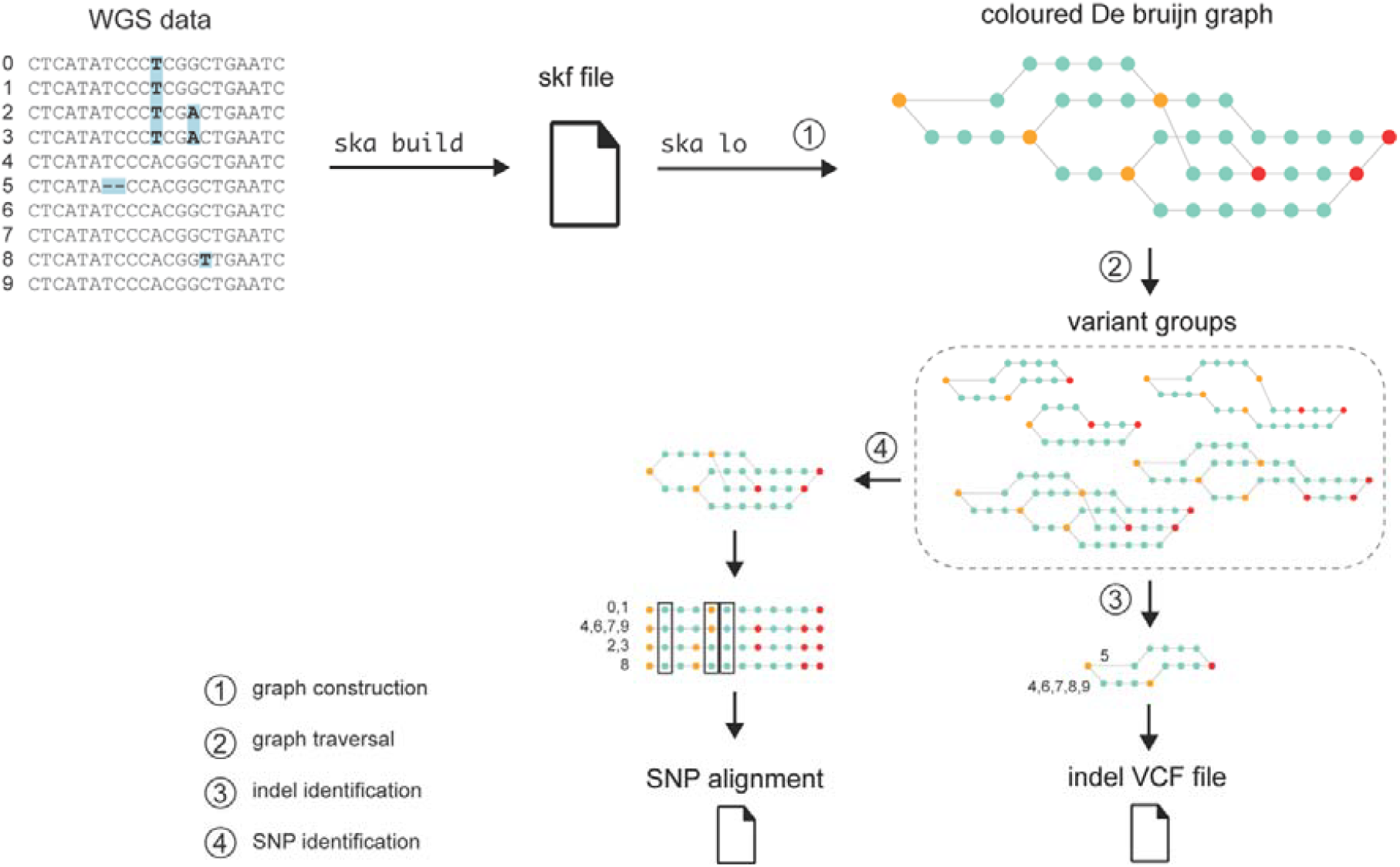
Schematic representation of ska lo analysis of WGS data. The circled numbers correspond to the algorithm’s steps as described in the main text, except for the optional step 5 (SNP positioning), which is omitted due to space constraints. The numbering next to the sequences represents their names. The graph’s orange and red nodes represent entry and exit nodes, respectively. In this example, the WGS dataset consisted of 10 samples, containing three SNPs and one indel, with split k-mers extracted using k = 7. ska lo inferred three SNPs (highlighted by rectangles in the path pseudo-alignment), while the indel was detected but not inferred as its fraction of missing data was 0.4.

Step 1: ska lo converts each split k-mer into two overlapping k-mers of size k-1, considering both forward and reverse-complement sequences. The combination of all overlapping k-mers constitutes a coloured De Bruijn graph in which nodes are k-mers of size k-1 and edges, which connect pairs of consecutive k-mers, are labelled (coloured) by the sample sets in which the two consecutive (k-1)-mers are found. ska lo then identifies entry nodes (nodes leading to two or more paths whose edges contain non-identical sample sets) and exit nodes (their reverse-complements) in the graph; the portion of the graph between an entry node and its corresponding exit node is referred to as a bubble. The graph is finally compacted by shortening paths between entry and exit nodes.

Step 2: the graph is traversed from each entry node using a recursive function taking as parameter a maximal recursive depth (d=4 by default). A variant group is then identified when at least two paths start from the same entry node and end at the same exit node. These can represent single graph bubbles of SNPs or indels, or correspond to multiple interconnected bubbles being traversed, generating a combination of variants. In high variant density regions, large variant groups would theoretically represent all variant combinations observed in the sample set.

Step 3: ska lo infers indels from variant groups having two paths and one of these paths corresponding to at most 2(k-1) nucleotides. Since indels are represented in both forward and reverse-complement orientations in the graph, they need to be deduplicated. Additionally, different insert sizes can be inferred for a given indel. Under the assumption that the shortest inserts are correct, these variant groups are sorted by the combined length of their paths and processed in increasing order. An indel is then validated if none of its surrounding k-mers have been found in an already validated indel, it corresponds to a non-ambiguous indel (ie, both variants are uniquely present in at least one sample) and its proportion of missing samples is below a threshold ‘m’ (m=0.1 by default). A variant call format (VCF) file is finally created, containing the different insert variants and their distribution among samples.

Step 4: ska lo then infers SNPs from each variant group by collecting the nucleotide column following each entry node in the pseudo-alignment of its paths. To ensure that paths are correctly aligned, variant groups are filtered to keep only paths of the most common length, which removes those containing an indel. However, paths of the same length can still be misaligned if one includes both an insertion and a deletion of the same size. To address this, ska lo discards paths containing at least ‘n’ indel surrounding k-mers recorded at the previous step (n=2 by default). As with indels, SNPs are represented in forward and reverse-complement in the graph, so need to be deduplicated. A given SNP might also be present in different variant groups, particularly in high-density variant regions, since variant groups are identified from each entry node. Since ska lo seeks to recover SNPs with their most complete set of samples, variant groups are sorted by the ratio of the number of paths to their length and processed in decreasing order. A SNP is then validated if it meets the same criteria as for indels: none of its surrounding k-mers have been found in an already validated SNP, it does not solely differ by an ambiguous nucleotide, and its proportion of missing samples is below ‘m’. Finally, ska lo outputs a SNP alignment for this sample set.

Step 5 (optional): if the user provides a reference genome, ska lo maps SNPs onto it, generating both a VCF file and a pseudo-genome alignment. Briefly, ska lo attempts to position variant groups on the reference genome and infers SNP locations accordingly (see Supplementary Methods for details). Variant groups that cannot be mapped are discarded, and their SNPs are excluded from the SNP alignment, VCF file, and pseudo-genome.

### Limitations of the algorithm

First, the removal of paths containing an indel can lead to ska lo missing SNPs. Any SNP located within k-1 nucleotides of an indel will always be missed since these will necessarily be located on the same path. In addition, the presence of an indel in a variant group will increase the fraction of missing samples of its SNPs since the sample set of the indel path is discarded. The other source of missed SNPs are MNPs. Since SNPs are identified among filtered variant groups through the presence of entry nodes, ska lo is only able to identify the first SNP and last SNP (in reverse-complement) of each MNP, meaning that internal SNPs, if any, are missed. There is, however, an exception to this: if an internal SNP is also present in another sample, its position will be ‘revealed’ and the SNP inferred (Supplementary Figure 1).

Second, ska lo can also produce erroneous calls. Its SNP-calling relies on the assumption that entry nodes among paths of equal length all correspond to SNPs. However, alternative variants can also preserve path lengths, such as micro-inversions or two undetected indels (i.e., within k-1 nucleotides) of equal length and opposite effect (i.e. one insertion and one deletion). Notably, in such cases, ska lo generates only two spurious SNPs, following the same logic as for MNPs. Finally, in highly polymorphic regions, complex variant groups might be incompletely identified if the recursive function fails to identify all paths. In such a case, the variant groups composed of these orphan paths can produce independent SNPs if their surrounding k-mers have not yet been visited, thereby creating duplicate SNPs. However, one of the two duplicates will necessarily have a proportion of missing data of at most 0.5, and setting the parameter ‘m’ above this value will discard these spurious SNPs.

### In-silico 20-samples benchmark

We first simulated an outbreak phylogeny of 20 isolates using Transphylo v1.4.10 (Xavier Didelot et al. 2017) with S. pneumoniae simulation parameters provided in (Derelle et al. 2024). Branch lengths of the phylogenetic tree were scaled up by different factors to simulate larger divergences using the same topology. Genomic variants were then simulated using phastSim (De Maio et al. 2022) for each of these modified phylogenies starting from the ATCC 700669 genome (NC_011900.1) at the root. phastSim parameters were set to generate on average one short indel (1-10bp) every 10 SNPs and one long indel (200bp) every 100 SNPs. The numbers of SNPs varied from 265 to 234,585. For each 20-sample dataset (ie, each level of divergence), the expected SNP alignment was reconstructed based on the tabular output of phastSim using a custom Python script. Simulated genomes generated by phastSim were then analysed by variant callers, and their SNP alignments were compared to the corresponding expected SNP alignments to infer true and false positive SNPs using compareALI v0.2 (https://github.com/rderelle/compareALI). Accession number of the strain SP264 genome was NZ_CP155532.1. OrthoANI values, a variation of the average nucleotide identities (ANI) (Yoon et al. 2017), were calculated using the online tool https://www.ezbiocloud.net/tools/ani. The versions of variant callers were: Snippy v4.6.0 (https://github.com/tseemann/snippy), SKA v0.4 (Derelle et al. 2024), parsnp2 v2.0.6 (Kille et al. 2024). Unless specified, ska lo was run with a maximum proportion of missing data of 0.1 (m=0.1). The phylogenetic trees, phastSim command-line parameters, phastSim simulated files, expected SNP alignments, and variant caller outputs have been deposited in Zenodo (https://zenodo.org/records/14863014).

### In-silico Mtb indel benchmark

Each pair of genomes was composed of the original root genome and the same genome mutated with 100 insertions of random sequences (random lengths between 1 and 10 nucleotides or fixed lengths for long insertions) at random genomic positions, in 10 replicates. The following constraints were imposed on insertions: (i) to not be located within 100bp from another insertion and (ii), for insertions of more than 1 nucleotide, the nucleotides corresponding to the extremities of the inserted sequences should be different to the nucleotides located on the opposite flanking insertion site. The latter constraint allows a unique alignment of these inserted sequences, which greatly facilitates their identification from read-alignment and ska lo outputs, making precise benchmarking possible. True positive indels were identified using the following two criteria: the insertion should have an exact match to one of the inserted sequences (forward or reverse-complement) and the original genome should display the alternative variant (i.e., no insertion at a position not treated as missing data). The Python scripts used to run this benchmark, along with the original genome assemblies, are available in the Zenodo file mentioned above.

### Analyses of the *S. aureus* outbreak dataset

Accession numbers of sequencing data were retrieved from https://microreact.org/project/SKASaureusSKAalign. The accession number of the reference genome used in these analyses was NZ_CDLR01000001.1 (strain SASCBU26). SNP alignment and polymorphic positions inferred by Snippy were extracted from the ‘clean full’ genome alignment using a custom python script.

### Within-strain analyses of *S. pneumoniae* PMEN2

Accession numbers of sequencing data and years of collection were retrieved from the supplementary data files of (Nicholas J. Croucher et al. 2014). Accession numbers of the reference genomes were CP002176.1 (strain 670-6B) and NC_011900.1 (strain ATCC 700669). The detection and removal of recombination tracks was performed using Gubbins v3.3.1 (N. J. Croucher et al. 2015). We used IQ-TREE v2.3.6 (Minh et al. 2020) for additional phylogenetic inferences. Root-to-tip analyses were performed using BactDating v1.1 with tree rooting using its initRoot function (Xavier Didelot et al. 2018). The clustering information distances used in the multidimensional scaling analysis were calculated using TreeDist v2.9.1 (Smith 2022).

### Efficiency measurements

Runtime and memory consumption measurements were performed on a MacBook Air laptop equipped with a M2 processor and 8 GB of RAM memory.

## Results

### Performance of ska lo SNP-calling in simulated benchmarks

To assess the performance of ska lo SNP calling, we built an in-silico benchmark using the genome of *Streptococcus pneumoniae* ATCC 700669 and a fixed phylogeny to generate sets of 20 genomes with increasing number of simulated variants (SNPs, short and long indels). These sets of genomes were analysed using the genome-aligner parsnp2, the popular read-alignment pipeline Snippy, and the k-mer based methods ska align, ska map and ska lo - all using a split k-mer size of 41. SNP alignments were trimmed to only retain variants not differing solely by an ambiguous nucleotide and positions with a maximum of 10% of missing data, considering ambiguous nucleotides as missing. The precision of variant callers was found to be above 0.99 in all cases except for parsnp2 at low SNP-densities using SP264 genome as reference (all values are provided in Supplementary data).

When using the simulated genome ‘1’ as a reference for variant calling, parsnp2 showed maximal sensitivity at low SNP densities due to the use of full genome assemblies without rearrangement, but declined significantly at higher SNP densities (Figure 2A). Snippy maintained relatively constant sensitivity (∼0.97) across datasets, while ska map exhibited slightly lower sensitivity (∼0.94 at low SNP densities) due to its use of short k-mers and a sharp drop at medium SNP densities due to its inability to detect MNPs. Using a closely related genome (SP264 strain; OrthoANI of 99.95% with the ATCC 700669 genome) as the reference reduced the sensitivity of all reference-based methods (e.g., ∼0.95 for Snippy), highlighting the impact of reference genome selection on reference-based variant calling.

**Figure 2:**
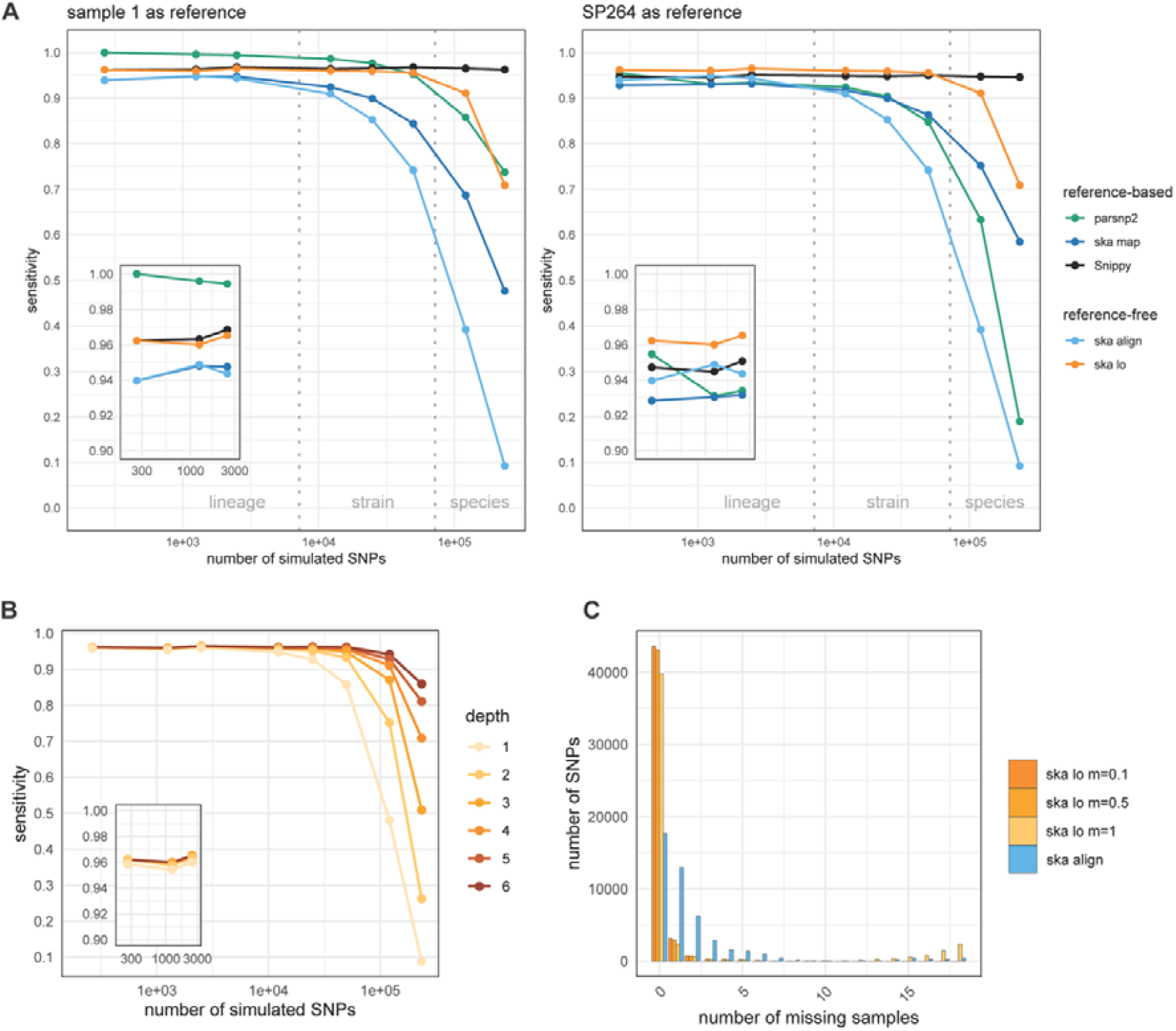
Benchmarking of SNP-callers. **A:** Sensitivities observed using two different reference genomes for reference-based callers. The vertical dotted lines marking the lineage-strain and strain-species boundaries correspond to average nucleotide diversities of 0.0005 and 0.005 respectively. Please note the log-scale on the x-axes, and that values obtained for the reference-free callers (ska align and ska lo) are identical in these two plots. Insets show sensitivities at lowest SNP-densities. **B:** Variation of ska lo maximum recursive depth parameter in benchmark analyses (default value is d=4). Inset shows sensitivities at lowest SNP-densities. **C:** Distribution of missing samples per SNP observed with the benchmark dataset of 49,732 simulated SNPs. ska lo was run with a maximum proportion of missing data ‘m’ of 0.1 (as in Figure 2 and 3B), 0.5 and 1, and ska align with m=1. In this dataset, ground truth SNPs have 0 or 1 missing sample.

Among reference-free methods, ska align matched ska map sensitivity at low SNP densities but dropped more sharply at medium densities, limited by both MNPs and overlapping variants. In contrast, ska lo achieved higher sensitivity (∼0.96) at low and medium SNP densities. This advantage stems from ska lo’s larger detection range of unique sequences, required for the identification of SNPs, spanning 2k-1 nucleotides compared to the k nucleotides of split k-mers. At this k-mer size, ska lo sensitivity matches or exceeds that of Snippy depending on the reference genome. ska lo also retained higher sensitivity at higher SNP densities compared to the other SKA commands as it accommodates MNPs and overlapping variants.

Investigating the influence of ska lo parameters, the maximum recursive depth for the graph traversal has no impact on the detection of SNPs at low and medium SNP-densities but higher values can further enhance ska lo performance at high SNP densities (Figure 2B). Despite this, ska lo remains unsuitable for the analysis of highly divergent genomes (ie, at the species level), with sensitivity still lower than that of Snippy and high computational costs associated with large graphs and their extensive traversal. The distribution of missing samples per SNP in a high-density dataset illustrates how effectively ska lo, run with the default maximum fraction of missing samples (m=0.1) or m=0.5, accommodates MNPs and overlapping variants compared to ska align (Figure 2C). However, removing SNP filtering based on missing data (m=1) resulted in a large number of SNPs at high fractions of missing samples, some of which may represent duplicate SNPs (see ‘Limitations’ section). It also significantly lowered the number of SNPs without missing samples, indicating that the sorting of variant groups before SNP calling can be improved.

We finally evaluated ska lo performance with SNP positioning on a reference genome, in which non-positioned SNPs are discarded. Considering low and medium SNP densities, ska lo sensitivity remained mostly unchanged when using the simulated genome ‘1’ as reference but dropped to that of Snippy when using the SP264 genome as reference (Supplementary data). Finally, we used the original ATCC 700669 genome as reference to compare genomic positions of SNPs inferred by ska lo to the simulated ground truth. The percentage of SNPs correctly positioned gradually decreased with variant density, from 100% to 97.6% (Supplementary figure S2). While this level of accuracy seems adequate for analysing SNP distribution across the genome (e.g., detecting recombination events), it might be insufficient for the accurate identification of known SNPs at specific genomic positions.

### Performances of ska lo indel-calling in simulated benchmarks

Assessing the performance of ska lo in indel calling within the previous benchmark is challenging, as indels can be inferred differently depending on the nucleotides surrounding the insert site. Even when considering all indels detected by ska lo as true positives, the percentage of inferred indels relative to the total number of simulated indels declined significantly at medium variant densities, with no noticeable impact of the maximum recursive depth threshold (Figure 3A). This decline occurs because, unlike in ska lo SNP-calling, indels are excluded from variant groups, and their detection is therefore influenced by overlapping variants.

**Figure 3:**
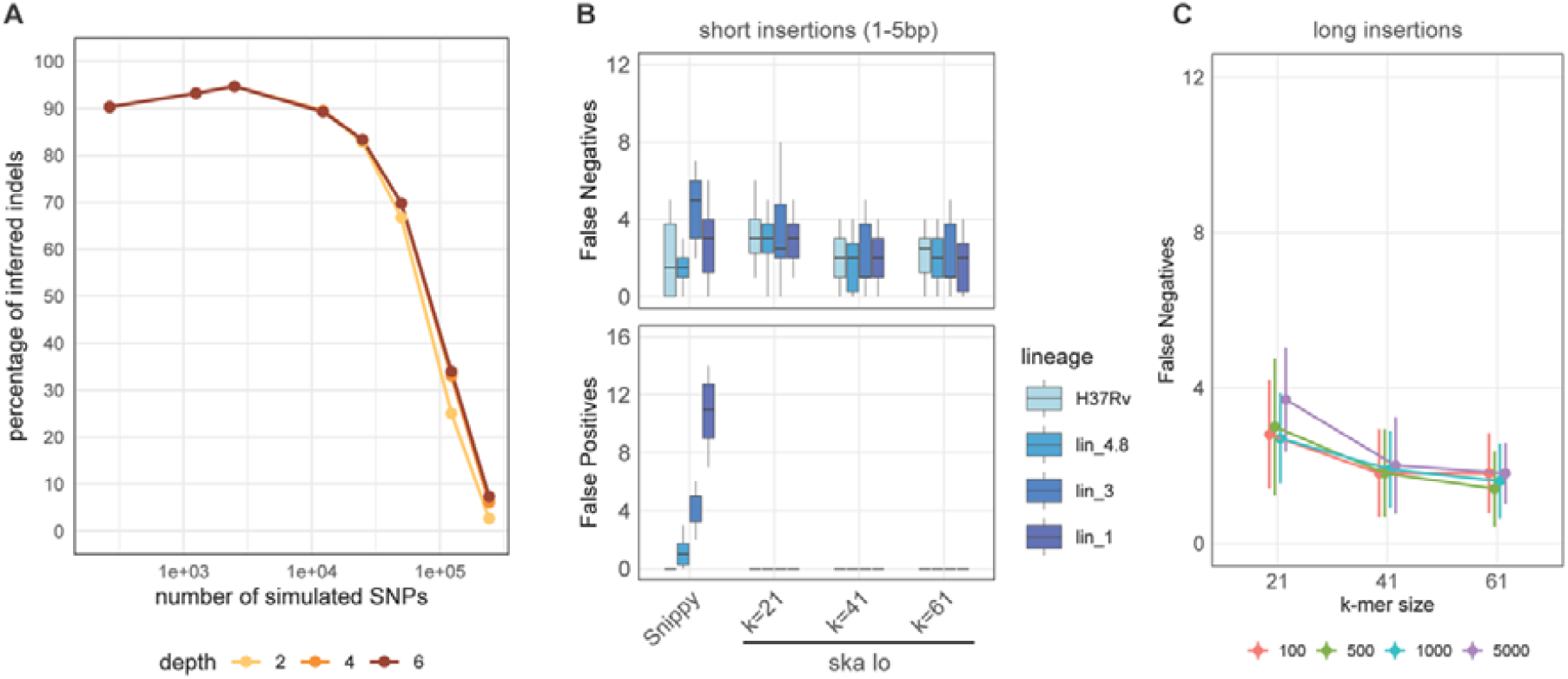
Benchmarkings of ska lo indel-calling. **A:** Percentage of inferred indels relative to the total number of simulated indels in the 20-isolate benchmark. The lines for maximum recursive depths of 4 and 6 are indistinguishable. **B:** False positive and false negative indels in the detection of short simulated insertions by Snippy and ska lo using genomes from different Mtb lineages. **C:** False negative indels in the detection of large simulated insertions by ska lo using the H37Rv genome. The different colours indicate different insert sizes, ranging from 100 to 5000 nucleotides. Error bars represent the standard deviations over 10 replicates

We then fully evaluated the accuracy of ska lo indel-calling in a low diversity setting by designing a second benchmark composed of pairs of Mycobacterium tuberculosis (Mtb) genomes differing by 100 randomly introduced synthetic insertions in 10 replicates. The insertion sites were selected to ensure that only a single possible indel could be inferred (see Methods), allowing us to identify false negative and false positive indels. These genome pairs were derived from assemblies representing different Mtb lineages, spanning increasing phylogenetic distances from the H37Rv reference genome, ranging from lineage 4.9 (strain H37Rv) to lineage 1, with sequence divergence below 0.001 SNPs per site. We compared ska lo indel inference using k-mer sizes of 21, 41 and 61 to that of Snippy.

ska lo missed on average 3 insertions out of 100 at k=21 and 2 insertions out of 100 at k=41 and k=61, a sensitivity level similar to that observed with Snippy (Figure 3B). Importantly, ska lo did not produce any false positive indel in any of these analyses, as opposed to Snippy which generated a number of false positives increasing with the genomic divergence to the reference genome. Accuracy to detect large indels, ranging from 100bp to 5kb, were only assessed for ska lo as these variants cannot be identified by standard read-alignment pipelines for which variant detection is limited by the read length (Ezewudo et al. 2018). ska lo was able to detect the longest insertions of 5kb, and displayed similar sensitivities to those observed with short insertions (Figure 3C). Three false positive indels were detected in all these analyses. However, manual inspection revealed that these actually corresponded to simulated insertions with longer than expected insert sizes, indicating that ska lo correctly identified the insertion events but failed to identify the shortest possible inserts.

### Variant calling in an outbreak setting

To evaluate how ska lo performs with real-world data, we first reanalysed a dataset of 55 *Staphylococcus aureus* samples from a study investigating an outbreak in a special care baby unit (S. R. Harris 2018; Coll et al. 2017; Simon R. Harris et al. 2013). Sequencing reads were analysed using Snippy, ska align, ska map, and ska lo, with k-mer sizes of 21 and 41. The reference genome used for reference-based calls corresponded to one of the 55 samples for which long-read sequencing had been performed. Only variants not differing solely by an ambiguous nucleotide and positions with a maximum of 10% of missing data were considered for the SKA commands. For Snippy, we used SNPs corresponding to the core-genome alignment and filtered indels inferred from each sample.

Snippy identified 92 SNPs in this dataset, ska align and ska map both produced 86 SNPs at both k-mer sizes, and ska lo identified 90 SNPs at k=21 and 92 SNPs at k=41 (Table 1). Running ska lo with SNP positioning on the reference genome produced the same results at both k-mer sizes, indicating that all variants were positioned. The comparison of VCF files produced by Snippy and ska lo at k=41 showed that 91 of the 92 SNPs were common to both tools. The single SNP identified only by Snippy was located next to a large unmapped genomic region and susceptible to misalignment errors (Supplementary Figure S3). The SNP exclusively detected by ska lo was shared by three samples forming a monophyletic group (position 1356572; samples “E” from (S. R. Harris 2018; Coll et al. 2017; Simon R. Harris et al. 2013)) and was inferred by Snippy but excluded from the core alignment due to the presence of missing data. As noted in (S. R. Harris 2018; Coll et al. 2017; Simon R. Harris et al. 2013), two MNPs were unexpectedly present in this low-diversity dataset—both composed of two SNPs separated by less than 10 nucleotides—accounting for the four-SNP difference between ska lo and the two other SKA commands at k=21. Increasing the k-mer size to 41 enabled ska lo to resolve more repeats, thanks to its larger detection range compared to split k-mers, and to identify two additional SNPs. Overall, these results align with the findings from the 20-isolate benchmark analyses at low SNP densities. Finally, Snippy inferred 18 indels across all samples, while ska lo inferred 13 and 19 indels at k-mer sizes 21 and 41, respectively. The comparison of the genomic sequences surrounding Snippy-identified indels and the sequences of bubble paths present in the ska lo indel vcf file at k=41 allowed us to identify 17 commonly inferred indels. Snippy inferred an additional 1bp insertion (position 50 in sample P14/B), while ska lo inferred two additional large deletions (55bp and 227bp, shared by samples HD and HT, and sample P25, respectively).

**Table 1:** Number of variants inferred from the S. aureus outbreak dataset by Snippy and the three SKA commands. For the SKA commands, the left and right values were obtained with k-mer sizes of 21 and 41 respectively.

|  | Snippy | ska align | ska map | ska lo |
| --- | --- | --- | --- | --- |
| number of SNPs | 92 | 86 / 86 | 86 / 86 | 90 / 92 |
| number of indels | 18 | – | – | 13 / 19 |

The analysis of the 55-samples skf file generated with k=41 took 3 seconds, 5 seconds, and 14 seconds for ska align, ska map, and ska lo respectively, with memory usages of 3.5 GB, 3.5 GB, and 1.7 GB. When using two CPUs, ska lo runtime decreased to 11 seconds. Factoring in the time required for skf file creation (8.4 minutes on 1 CPU), the runtime increase for ska lo remained relatively modest.

### Within-strain phylogenetic analyses

Phylogenetic reconstructions at the strain level usually involve the detection and removal of recombination events using a reference genome (N. J. Croucher et al. 2015; X. Didelot and Wilson 2015). For this set of analyses, we used 170 *Streptococcus pneumoniae* PMEN2 isolates, many of which represented a nationwide outbreak of multidrug-resistant disease in Iceland (Nicholas J. Croucher et al. 2014). Sequencing reads were analysed using Snippy, ska map and ska lo, both using a k-mer size of 21.

Using the *S. pneumoniae* 670-6B genome as reference genome, as in (Nicholas J. Croucher et al. 2014), Snippy and ska map identified 23,469 and 11,950 SNPs respectively. ska lo, using a maximum fraction of missing data per position of 0.4, identified 17,548 SNPs, of which 1,048 could not be positioned. Despite these differences, the detection of recombination tracts by Gubbins revealed similar patterns (Supplementary Figure S4), and produced similar numbers of filtered SNPs: 2,320, 2,234 and 2,280 filtered SNPs for Snippy, ska map and ska lo, respectively. Among these, 1,990 polymorphic positions were shared by all three variant callers (Supplementary figure S5). In the absence of an objective ground truth, we relied on the concordance of these filtered SNPs with the assumption of a molecular clock to assess their accuracy. Root-to-tip analyses based on Gubbins final trees and trees produced by IQ-TREE under the GTR model revealed ska lo dataset having the strongest temporal signal (R^2^= 0.66 in both cases; Figure 4A), followed by ska map (R^2^= 0.61 and 0.62), then Snippy (R^2^= 0.46 and 0.5; Supplementary figure S6). However, the estimated date of the most recent common ancestor (MRCA) for this group of isolates was similar across all three analyses. Removing the SNP filtering based on missing data (m=1) increased the number of SNPs identified by ska lo (20,093 SNPs of which 2,114 not positioned; 2,550 filtered SNPs after Gubbins analysis), but was associated with a notable weakening of the temporal signal (Supplementary figure S6). This result highlights again the presence of spurious SNPs generated by ska lo at high fractions of missing samples when analysing high SNP-density regions, and the need to keep the value for this parameter below 0.5.

**Figure 4:**
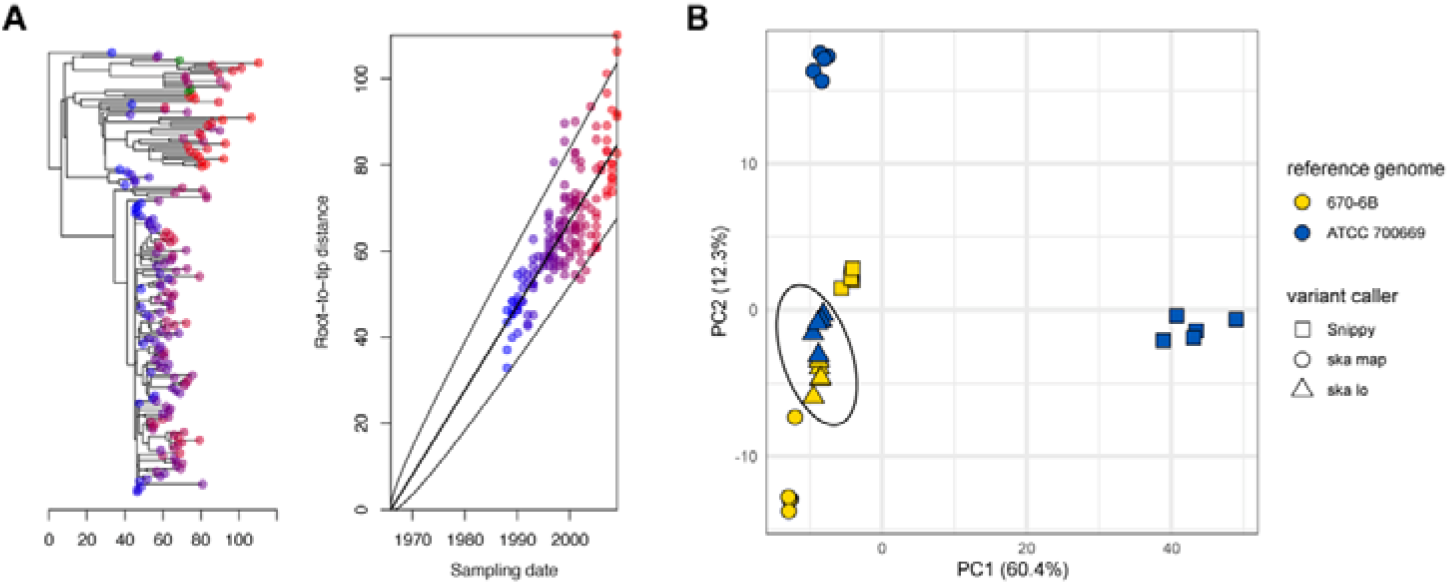
*S. pneumoniae* PMEN2 isolates analyses. **A:** BacDating root-to-tip analysis based on a tree obtained using IQ-TREE under the GTR model and Gubbins filtered SNP set produced by ska lo (genome 670-6B as reference). **B:** Multidimensional scaling analysis of phylogenetic trees based on clustering information distances (Smith 2021). For each variant caller and each reference genome, five trees were generated using IQ-TREE under the GTR model to account for the uncertainty in phylogenetic inferences. The dashed ellipse circles all trees based on ska lo filtered SNPs.

To test the practical limits of the variant callers, we repeated these analyses using a more distant reference genome (*S. pneumoniae* ATCC 700669; OrthoANI of 98.52% with the 670-6B genome). Here, the filtered SNP sets obtained after Gubbins analyses differed significantly among the three variant callers, with Snippy, ska map, and ska lo yielding 3,291, 2,082, and 2,040 filtered SNPs, respectively, and only 1,537 polymorphic positions in common (Supplementary Figure S7). The large number of filtered SNPs from Snippy was likely due to misalignment errors caused by the distant reference genome (Falconer et al. 2022; Yoshimura et al. 2019), which resulted in phylogenetic inferences that deviated greatly from those obtained using the 670-6B genome (Figure 4B) and weaker temporal signal (R^2^= 0.25 and 0.29; Supplementary figure S8). In contrast, alignment-free callers ska map and ska lo produced phylogenetic trees that were more consistent between the two sets of analyses and retained stronger temporal signals (R^2^= 0.45 and 0.58 for ska map and ska lo respectively). This observation was particularly evident for inferences based on ska lo SNPs, as the topologies from the two analyses were nearly identical. The minimal impact of the reference genome on ska lo-based phylogenetic analyses might be attributed to the fact that ska lo is also a reference-free caller that only subsequently maps SNPs to the reference genome in bulk, thereby preserving their relative distribution along the genome (average length of mapped variant groups of ∼150 bp; Supplementary Figure S9).

In terms of efficiency, ska lo took 34 seconds (27 seconds using 2 CPUs) and used 2.4 GB of memory to infer SNPs and indels from the 170-samples skf file and position SNPs on the 670-6B reference genome. In comparison, ska map took 17 seconds and required 7.6 GB of memory.

## Discussion

In this study, we introduced ska lo, a graph-based approach derived from split k-mer files to address some of the limitations of existing reference-free methods in detecting SNPs. ska lo proved effective in both benchmark and real-world data analyses, demonstrating its capability as a reference-free variant caller. Our results highlight once again the importance of reference genome choice in alignment-based variant calling. ska lo performs similarly to Snippy when the reference genome is closely related to the studied isolates, as seen in the *S. aureus* outbreak analysis. However, when the reference genome is more distantly related to them, as in the within-strain analyses of *S. pneumoniae*, its reference-free mode avoids alignment errors and reference bias. Finally, It is important to note that higher performance metrics do not necessarily translate into better results. In the analysis of the *S. aureus* outbreak, read-alignment and ska align SNP inferences led to phylogenies with identical topology (Harris, 2018), and all root-to-tip analyses based on the *S. pneumoniae* dataset produced similar MCRA dates. It remains to be determined whether the improvement in sensitivity provided by ska lo can lead to better biological or epidemiological insights, with existing SKA commands already performing well as demonstrated by recent studies (Mäklin et al. 2022; Chew et al. 2023; Maechler et al. 2023).

We identified three practical limitations of ska lo. First and foremost, its usage should be limited to within-strain analyses and lower diversity datasets. At higher divergences, ska lo simplistic logic struggles with high variant densities, resulting in relatively low sensitivity. Notably, our benchmark was particularly unrealistic at high SNP densities, as it only accounted for SNPs and indels as sources of genomic variation. If gene duplications, lateral gene transfers, and recombination events were included (ref dynamic genome bacteria), its sensitivity drop might be even more pronounced. For these reasons, we recommend partitioning WGS datasets into low-genomic-diversity groups, such as lineages, strains of serotypes, using MLST schemes (Uelze et al. 2020) or PopPunk (Lees et al. 2019). Please note that a pipeline integrating SKA and PopPunk has recently been proposed to facilitate these steps (McHugh et al. 2024). Second, ska lo produces spurious SNPs at high fractions of missing samples when highly polymorphic regions are analysed. While this artifact can be mitigated by setting the maximum fraction of missing data parameter below 0.5, it also limits confident inference to SNPs in the core genome. Finally, the accuracy of ska lo SNP positioning on a reference genome decreases with variant density. Without further validation, we recommend against using ska lo SNP genomic positions to test for the presence or absence of specific variants (e.g., SNPs causing antibiotic resistance in Mycobacterium tuberculosis (The, CRyPTIC Consortium 2022)).

In line with the SKA toolkit, ska lo offers a fast and simple-to-use solution for reference-free variant calling. The main parameter controlling its behaviour, the maximum recursive depth for graph traversal, has been set sufficiently high to handle any WGS dataset for which the tool was designed out of the box. Its development reflects an ongoing wave of simple, fast, and accurate k-mer-based programs designed to facilitate and democratize bioinformatics analyses (Bray et al. 2016; Lu et al. 2022; Lees et al. 2019).

## Conflict of Interest

The authors declare that there are no conflicts of interest.

## Acknowledgements

RD, KM and AL were supported by the NIHR Health Protection Research Unit in Respiratory Infections, in partnership with the UK Health Security Agency [NIHR200927]. JAL, JH and VRB were supported by the European Molecular Biology Laboratory. NC. and LC acknowledge funding from the MRC Centre for Global Infectious Disease Analysis (reference MR/R015600/1), jointly funded by the UK Medical Research Council (MRC) and the UK Foreign, Commonwealth × Development Office (FCDO), under the MRC/FCDO Concordat agreement and are also part of the EDCTP2 programme supported by the European Union.

## Notes

### Competing Interest Statement

The authors have declared no competing interest.

### Summary of Updates

skalo has been integrated into SKA2 (command: ska lo) and now also infers SNPs, which is the main focus of this revised version.

