## Supplementary figures for "Reference-free variant calling with local graph construction with ska lo (SKA)"

**Figure S1**

The number of SNPs inferred by skalo for each of these two examples (using k=7) are indicated on the right.

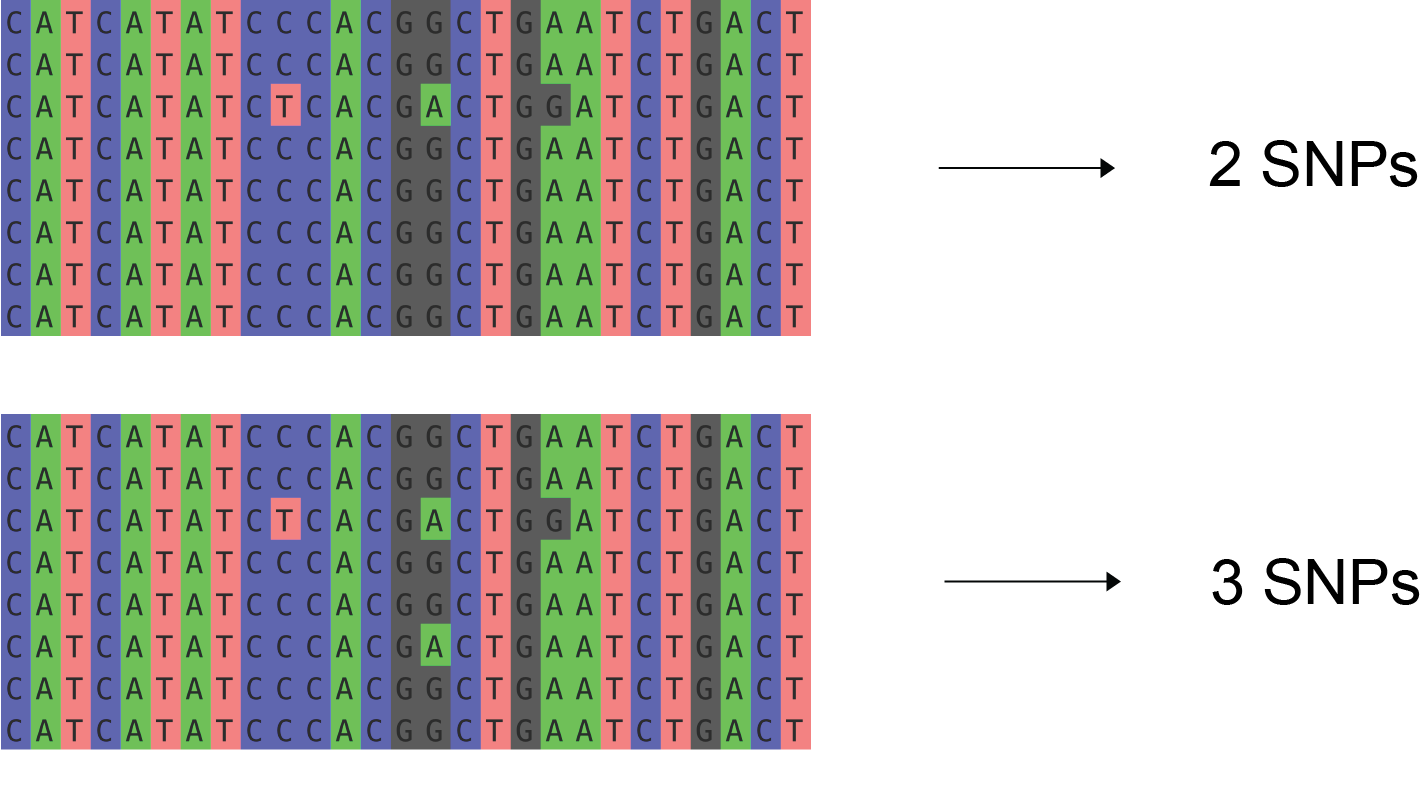

**Figure S2**

skalo SNP positioning in benchmark analyses (ATCC 700669 genome as reference). The plot shows the proportions of mis-positioned SNPs as percentages of the total number of positioned SNPs.

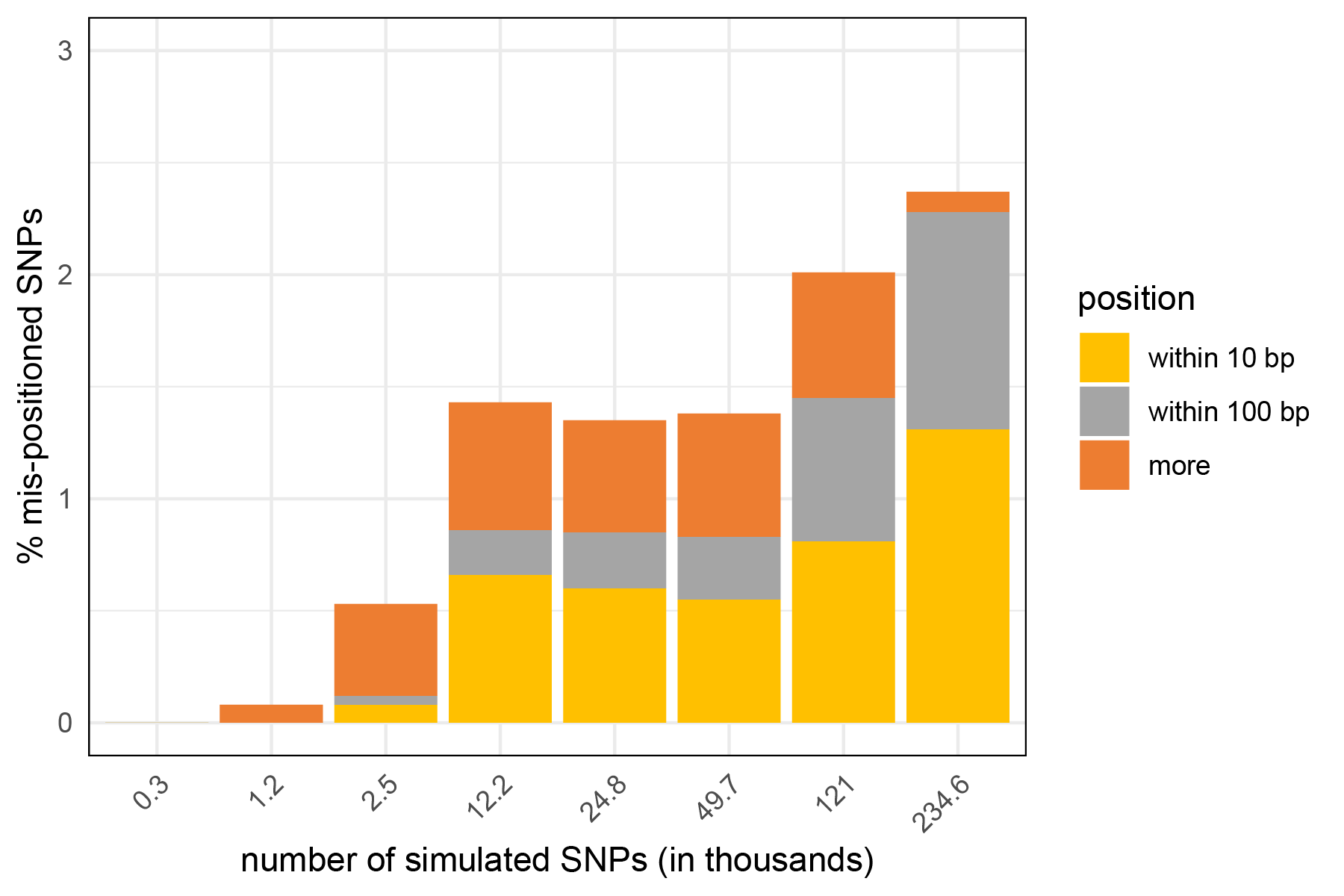

**Figure S3**

Visualisation using the command ‘samtools tview’ of the SNP specific to Snippy in the analysis of the *S. aureus* outbreak: sample P25 (ERR128708) had a mutation at position 1186254 that potentially corresponded to read misalignment around an indel:

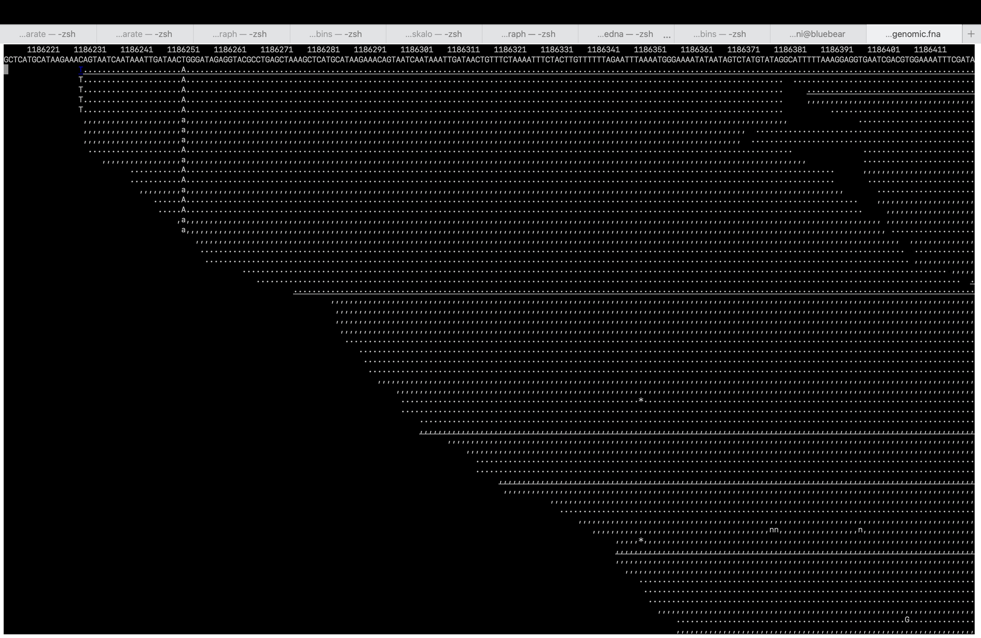

**Figure S4**

Phandango visualisation (<https://jameshadfield.github.io/phandango/#/main>) of Gubbins outputs obtained using the 670-6B genome as reference.

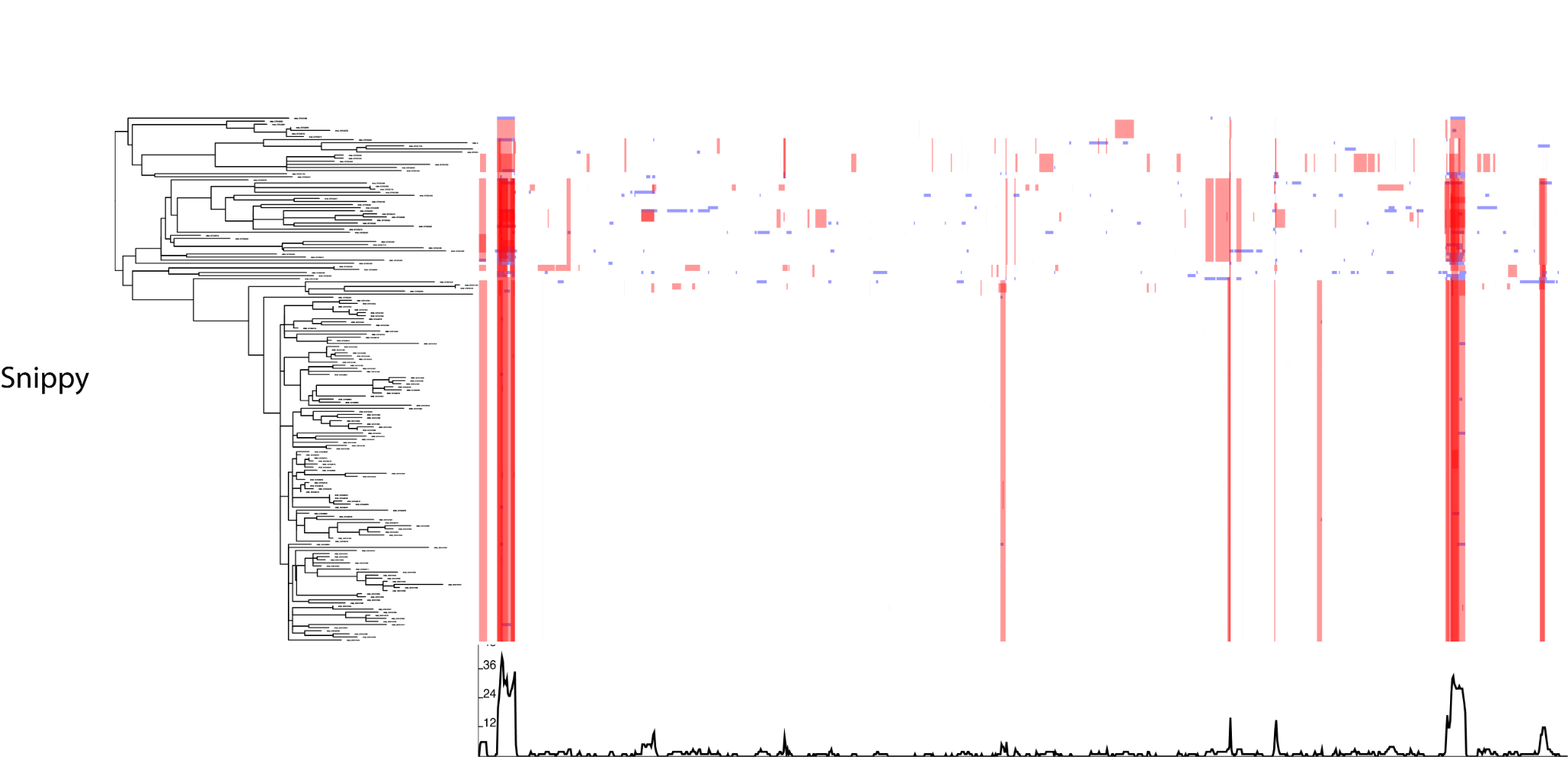

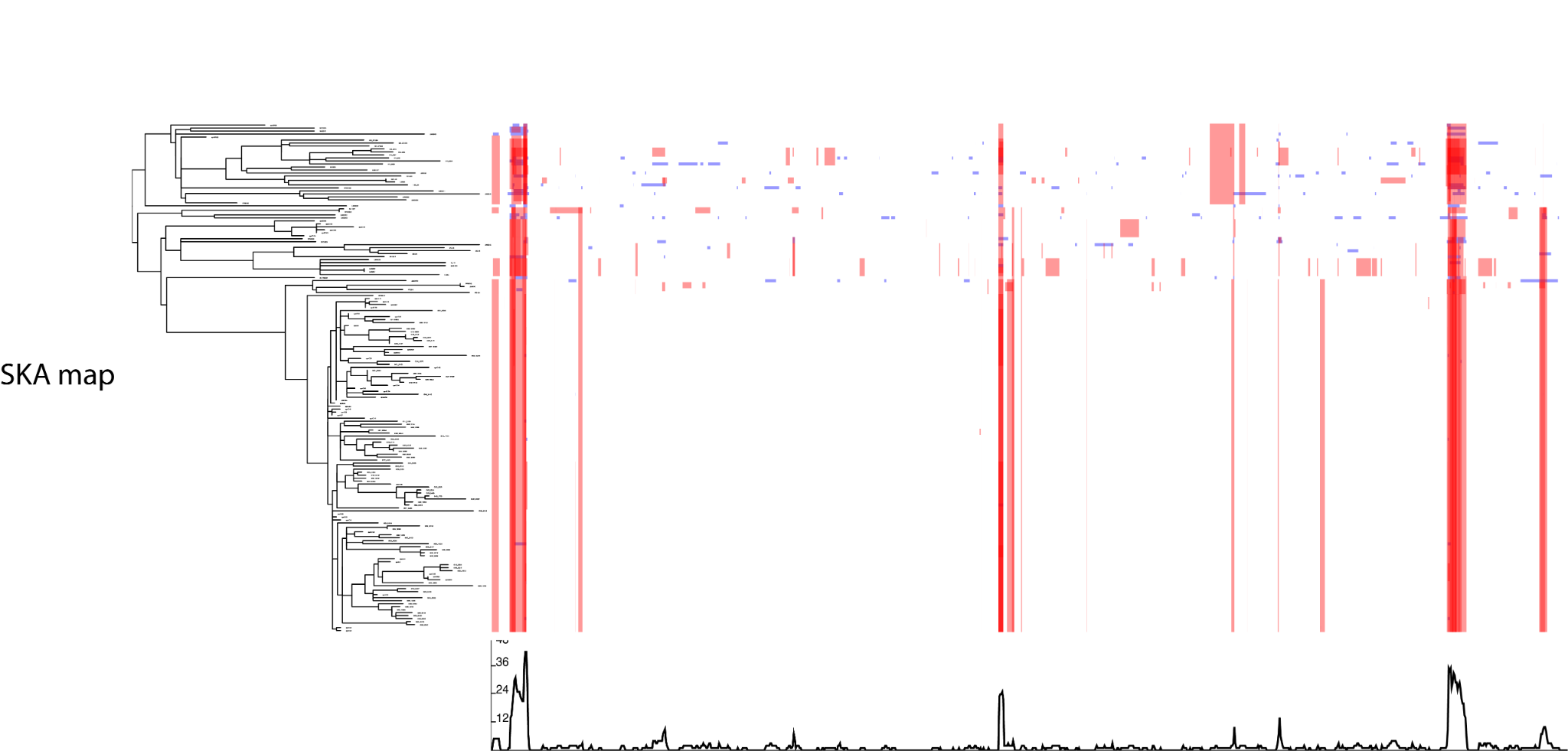

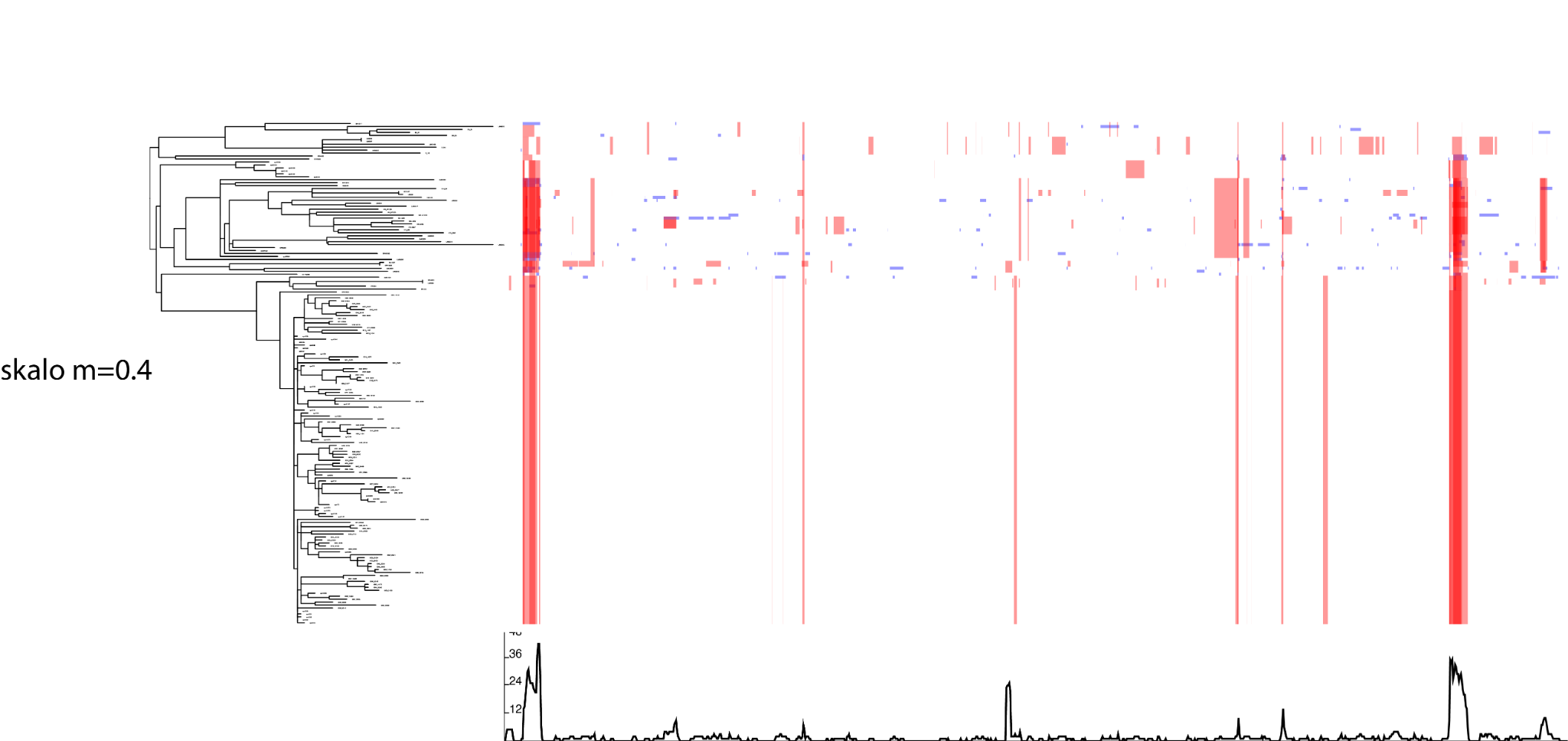

**Figure S5**

Venn diagram based on 670-6B genomic positions of Gubbins filtered SNPs sets obtained from the three variant callers.

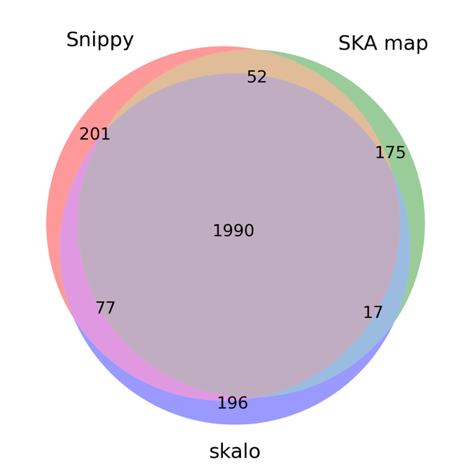

**Figure S6**

BacDating root-to-tip analyses of the 670-6B reference genome analyses. For each variant caller, the left and right plots are based on the Gubbin final trees (RaXML under the GTRGAMMA model) and trees obtained from the filtered sets of SNPs using IQTREE under the GTR model respectively.

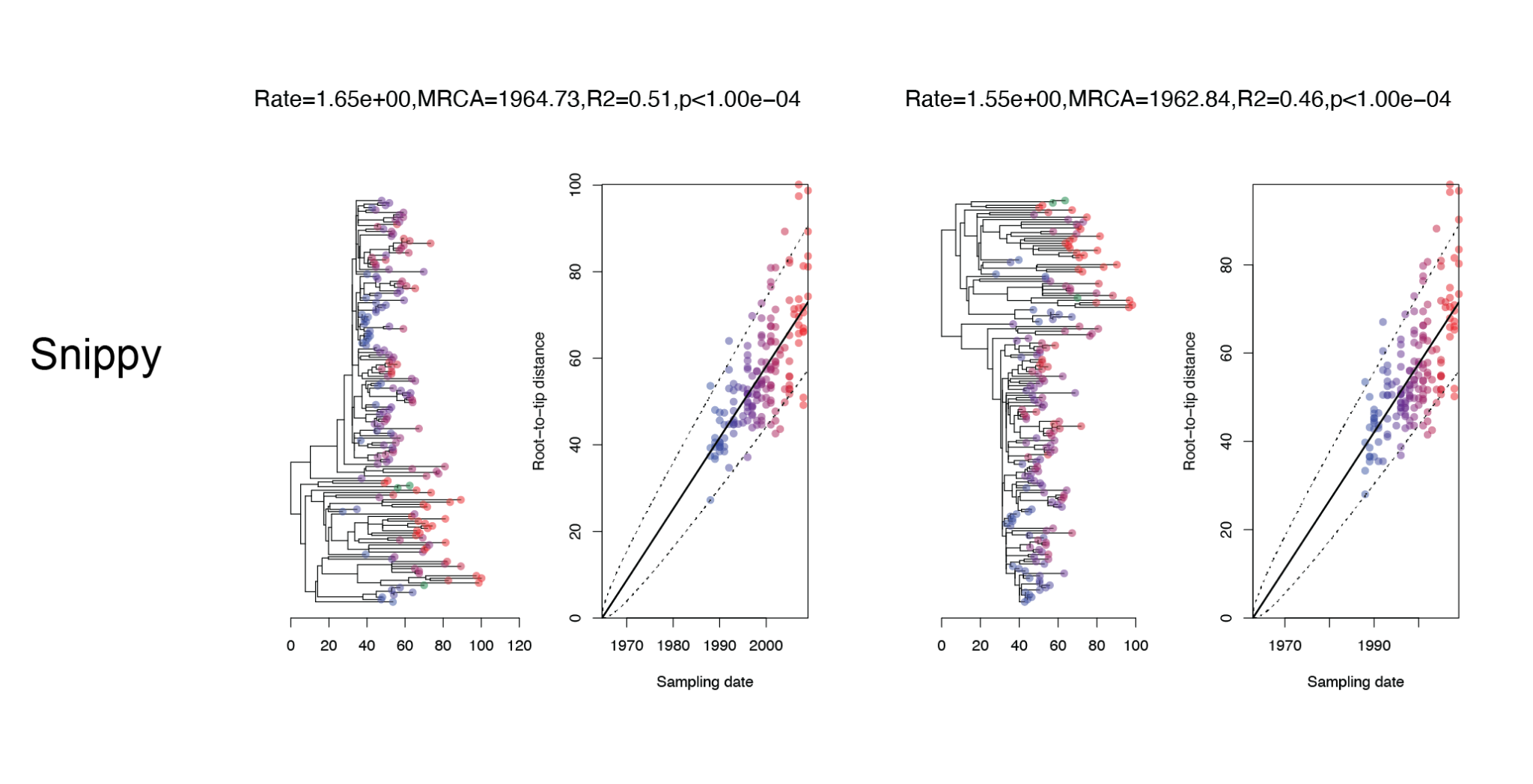

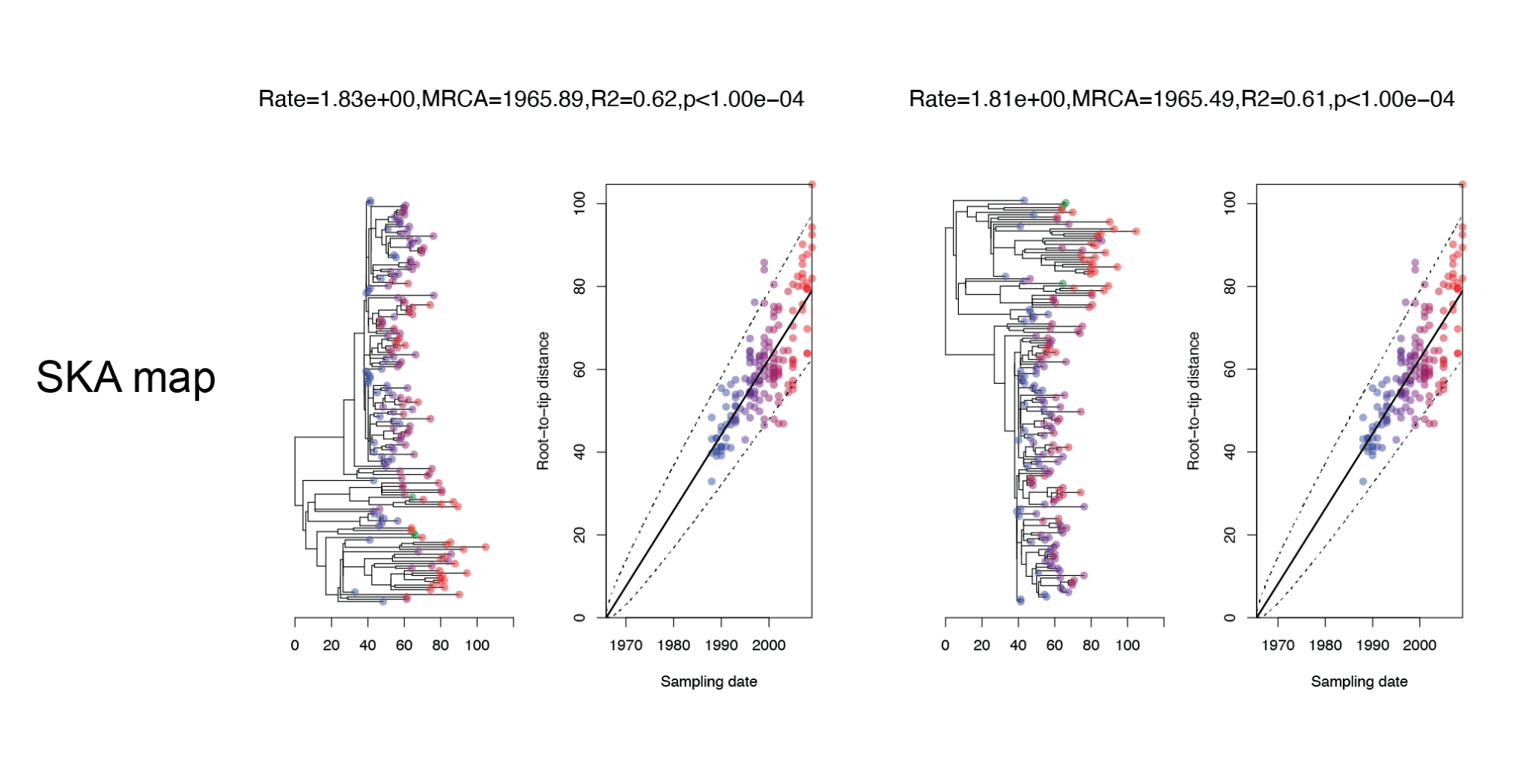

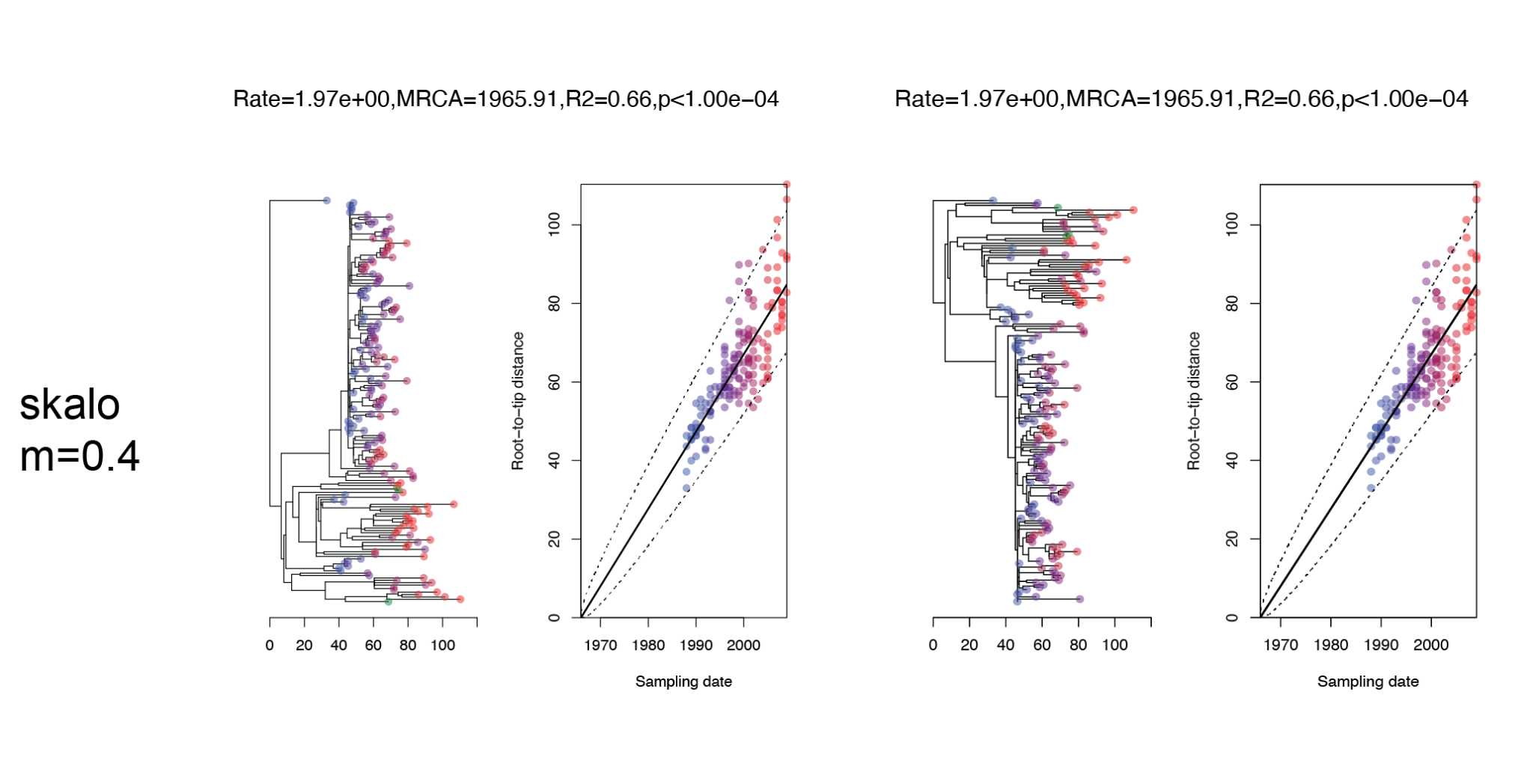

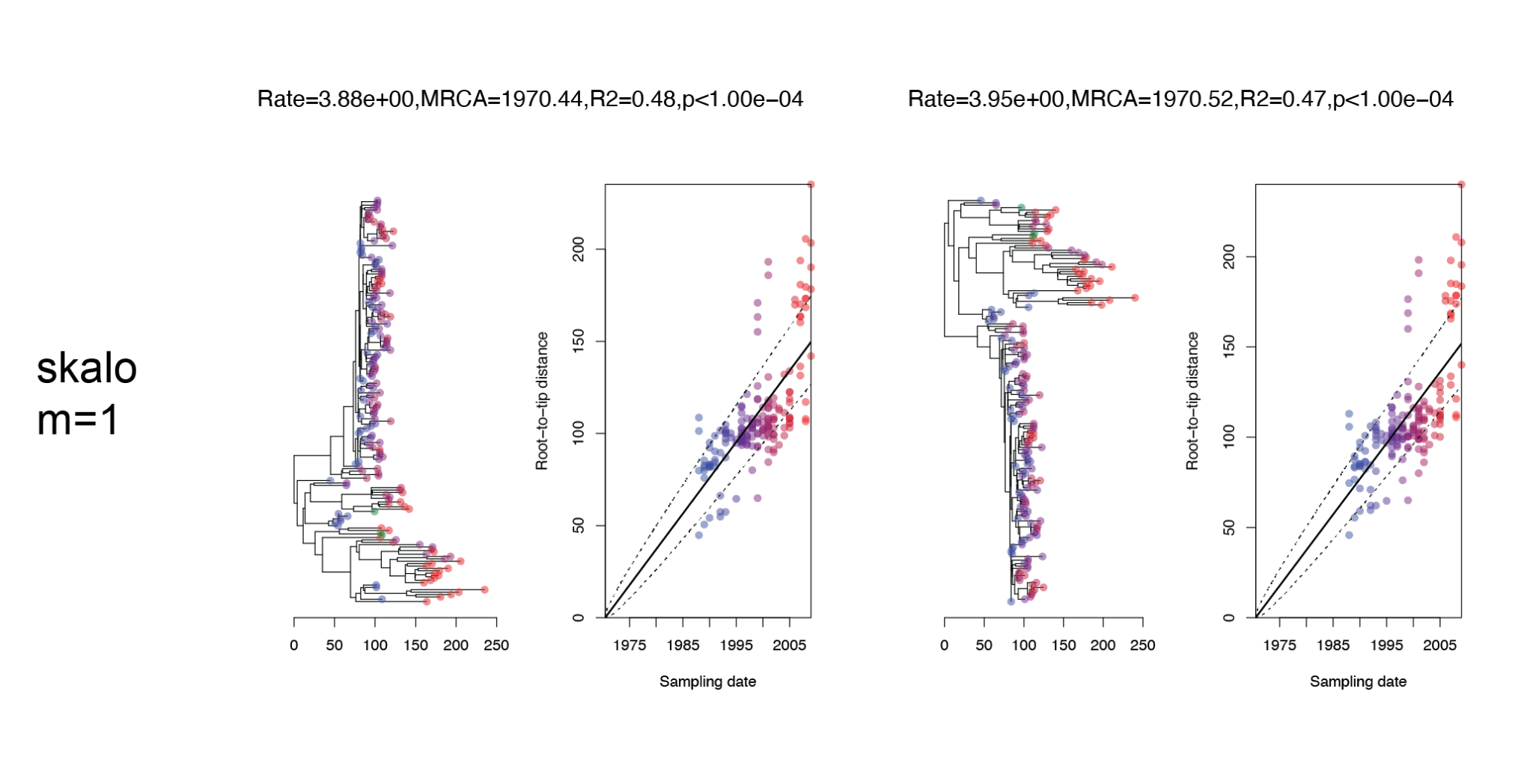

**Figure S7**

Venn diagram based on ATCC 700669 genomic positions of Gubbins filtered SNPs sets obtained from the three variant callers.

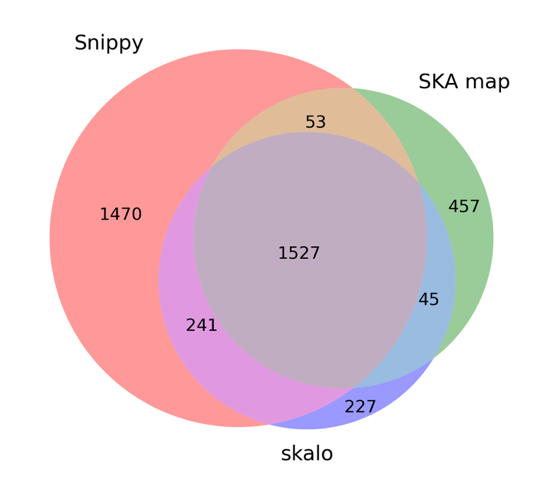

**Figure S8**

BacDating root-to-tip analyses of the ATCC 700669 reference genome analyses. For each variant caller, the left and right plots are based on the Gubbin final trees (RaXML under the GTRGAMMA model) and trees obtained from the filtered sets of SNPs using IQTREE under the GTR model respectively.

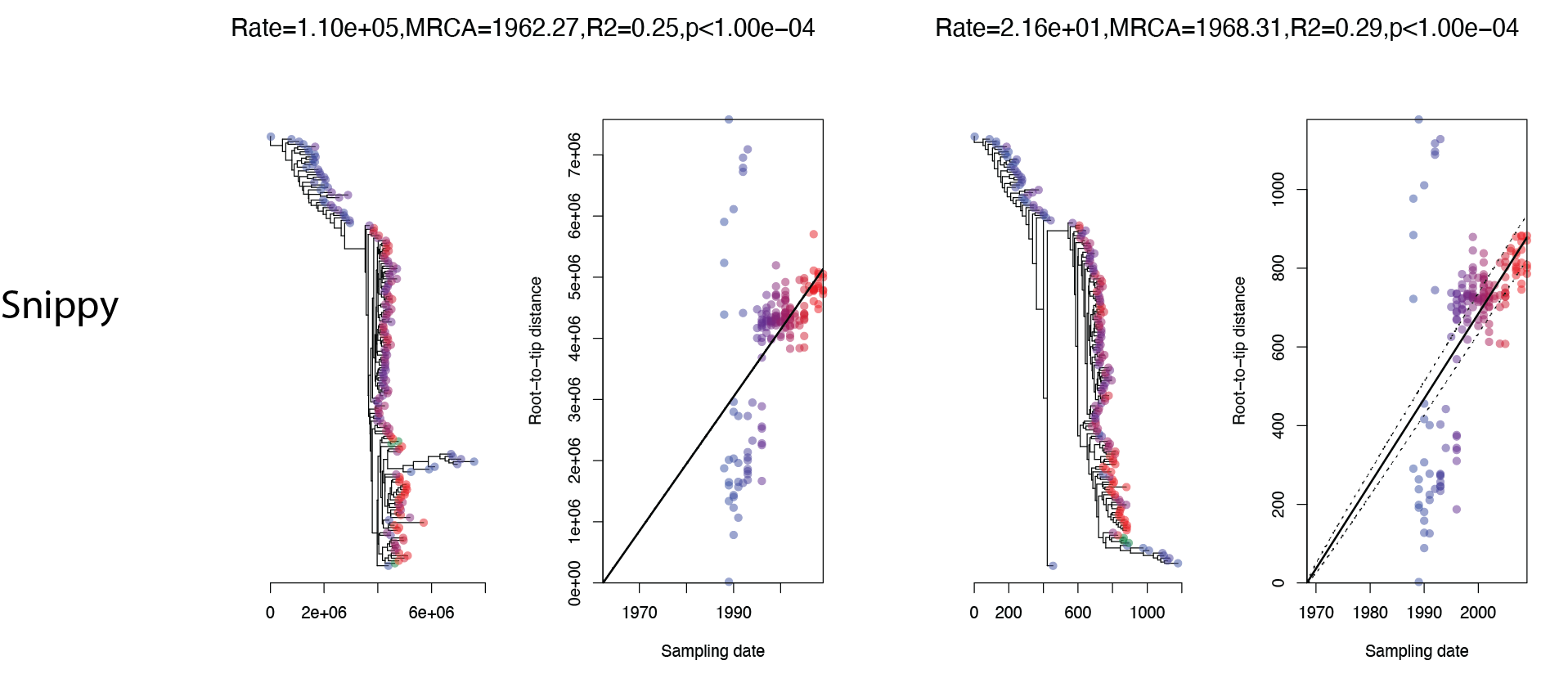

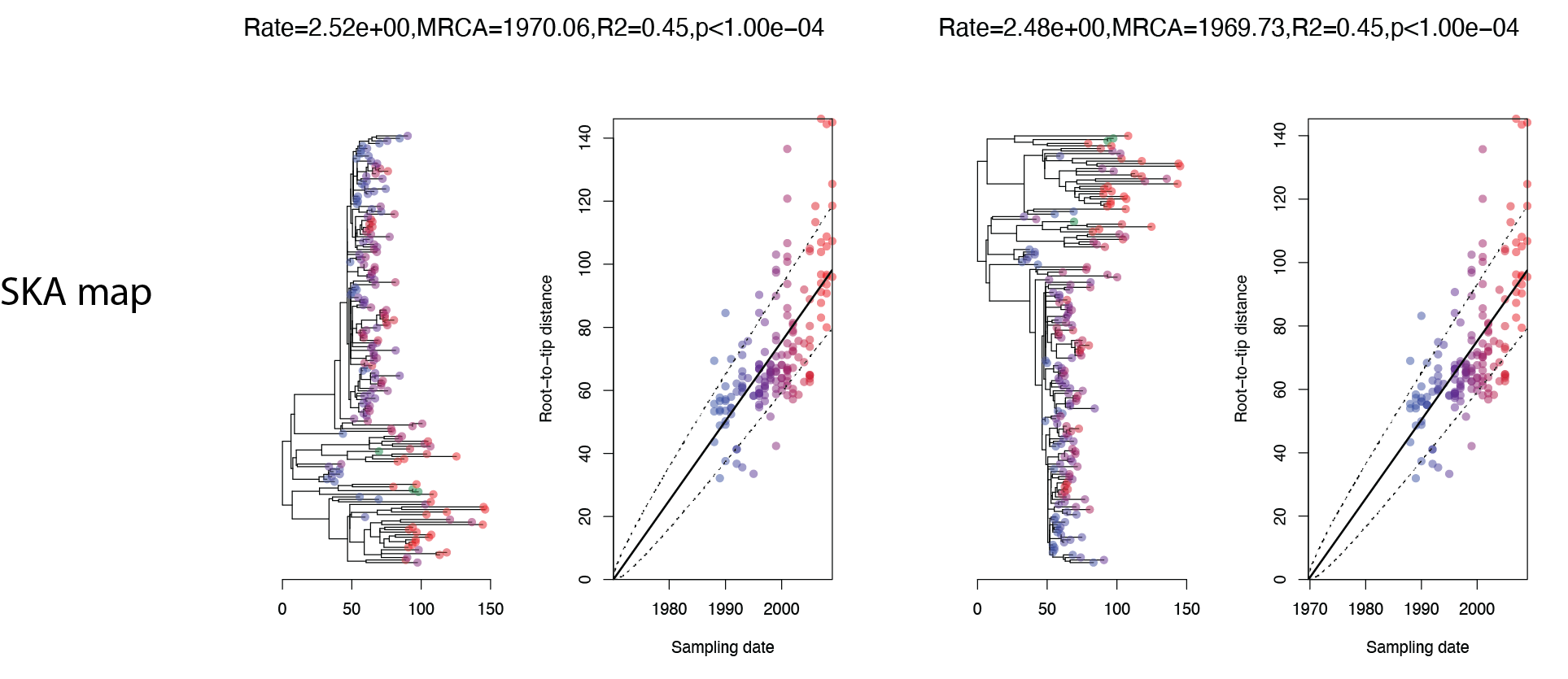

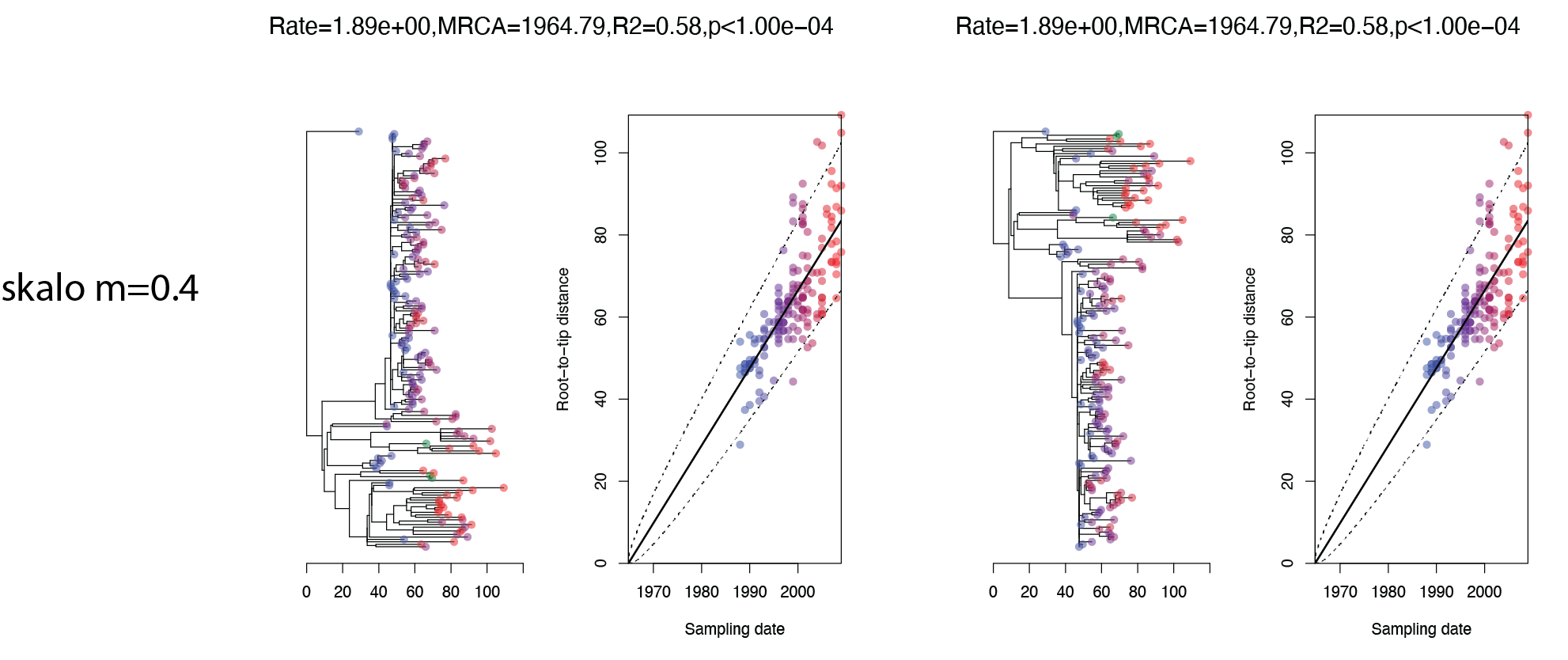

**Figure S9**

Distributions of the length of variant groups in bp when SNPs were positioned onto the reference genomes in the Spe PMEN2 analyses (one variant group length per positioned SNP). The lower bond of the distributions corresponds to single isolated SNPs: 40 bp as the dataset was analysed using k=21. The averages of the distributions were 146 and 151 bp for the 670-6B and ATCC 700669 reference genomes respectively.

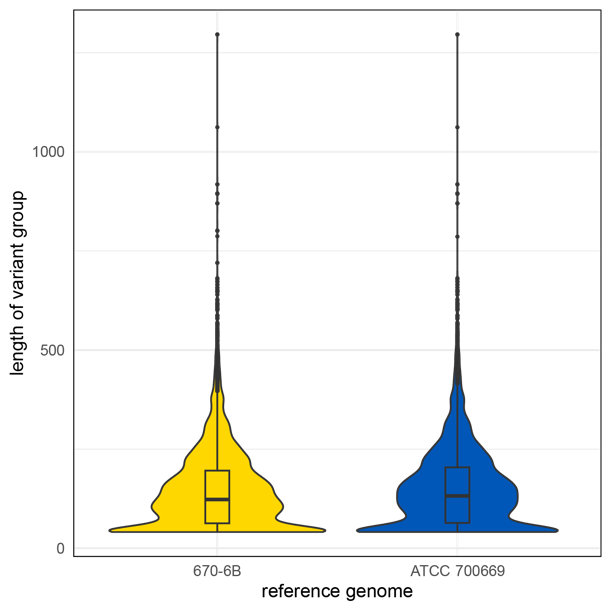
