## Supplementary methods for "Reference-free variant calling with local graph construction with ska lo (SKA)"

SNP positioning on a reference genome

The current implementation of SNP positioning on a reference genome contains two hardcoded parameters (highlighted in red in the description below). These parameters will be either removed or implemented as proper arguments in the ska lo command in future versions.

If a reference genome is provided by the user, it is fragmented into (k-1)-mers (hereafter referred to as ‘k-mers’ for simplicity) in both forward and reverse-complement orientations. K-mers, along with their positions and orientations, are stored if they appear no more than three times in the reference genome.

Variant groups containing at least one validated SNP are then considered for positioning. ska lo assigns a genomic position to each variant group's entry node based on the most frequent position of all its k-mers in the reference k-mer map. For each k-mer, if a match is found, the corresponding position and orientation in the reference genome are recorded, adjusting for the k-mer offset. The most frequent combination of position and orientation among all observed matches is selected as the final variant group position, ignoring those with fewer than ten occurrences to avoid incorrect placements. In the case of a tie (multiple positions occurring with equal frequency), the variant group is not positioned.

If a genomic position is determined for a variant group, individual SNPs are positioned on the reference genome based on the variant group's position, orientation, and their relative locations within the group. Conversely, variant groups that cannot be positioned are discarded.
